# Genetic combination risk for schizophrenia

**DOI:** 10.1101/478958

**Authors:** Kengo Oishi, Tomihisa Niitsu, Nobuhisa Kanahara, Tasuku Hashimoto, Hideki Komatsu, Tsuyoshi Sasaki, Masayuki Takase, Yasunori Sato, Masaomi Iyo

## Abstract

Schizophrenia is a highly hereditary mental disease^1^ related to abnormal dopaminergic activities.^2,3^ To elucidate the mechanisms underlying schizophrenia’s development, genomic studies have sought to identify the pathogenic genetic polymorphisms. Large-scale genome-wide association studies (GWAS) have reported potential candidate loci that contribute to schizophrenia’s development.^4,5^ The risk genetic profiles are not yet established. Here we show that the combination of three functional single nucleotide polymorphisms (SNPs) related to the key factors of dopaminergic signaling can be used to predict the risk of schizophrenia’s development, though none of the SNPs is known to be associated by itself. These functional SNPs were reported to demonstrate directional influences in their parent gene activity, perhaps characterizing the integrated properties of dopaminergic signaling. Interestingly, the risk combination presented here included the major genotype as well as the minor polymorphisms, suggesting a possible association of unaffected activities of some dopamine-related genes with the disease development. The phenotype speculated based on the allelic status seemed consistent with the conventional pathophysiological hypotheses, although recently developed predictive methods, such as the polygenic risk score, could miss this potent pathogenic role of carrying a normal genotype by evaluating only minor polymorphisms. Our results demonstrate the presence of a subtype in schizophrenia with the favored genetic background related to dopamine signaling. Our findings indicate the possibility that the combinations could characterize integrated biological functions (including neurotransmission) and therefore identify individuals with a disease risk. The biological microenvironment indicated by the functional SNPs could bring an insight to elucidate the pathogenic mechanisms of developing schizophrenia. Furthermore, we believe that our approach will contribute to the development of innovative means to predict disease risks even for other multi-factorial diseases and then, the following preventive medicine.

## Main

Schizophrenia is a disabling chronic mental illness, consisting of the prodromal phase and following psychotic onset, with the lifetime prevalence of 0.4% to 1.0% worldwide.^6,7^ The majority of first-episode schizophrenia patients positively respond to antipsychotic treatment with dopamine D2 receptor (DRD2) antagonists.^2^ Increased dopamine synthesis was observed in these responding patients,^3^ suggesting the involvement of abnormal dopaminergic signaling in the pathology of schizophrenia. Although epidemiologic studies convincingly suggested the high genetic heritability of schizophrenia,^1^ genome-wide association studies (GWAS) have not succeeded in indicating the apparent association of dopamine-related genes with schizophrenia.^8^ In fact, those studies tended to obtain negative results regarding the genetic associations with schizophrenia among the candidates proposed by a variety of psycho-pharmacological hypotheses.^9^

We recently demonstrated that analyses of the combinations of multiple functional single nucleotide polymorphisms (SNPs) of the parent genes that are key factors in dopaminergic signaling could be used to predict the risk of developing late-onset treatment-resistant schizophrenia (L-TRS).^10^ L-TRS may be due to antipsychotic-induced dopamine supersensitivity;^11,12^ i.e., L-TRS may be caused by the compensatory up-regulation of DRD2.^13–15^ Because individuals with L-TRS may be correctly classified as having typical dopamine-related schizophrenia, we evaluated the combination of dopamine-related functional SNPs, i.e., rs10770141 of the tyrosine hydroxylase (*TH*) gene, rs4680 of the catechol-O-methyltransferase (*COMT*) gene, and rs1800497, also called TaqIA, of the *DRD2* gene (Table 1). We identified two potent risk combinations for L-TRS; one is the double SNP combination of T(+) of rs10770141 and Met(−) of rs4680, and the other is the triple SNP combination of T(+) of rs10770141, Met(−) of rs4680, and A1(+) of rs1800497. It has been reported that these individual SNPs may reflect the relatively higher *TH* gene expression^16^ and COMT activity.^17^ In addition, the individuals with A1(+) of rs1800497 could have lower *DRD2* gene expression.^18^ Since these dopaminergic characteristics could be associated with schizophrenia, we tested here whether these allelic combinations could pose a risk for the development of the disease.

**Table 1.** The functional SNPs

| SNP ID | Related gene | Polymorphism function [ref.] | WT | Poly-morphism | MAF |
| --- | --- | --- | --- | --- | --- |
| rs10770141 | *TH* | 30%–40% higher gene expression [16] | C | T | 6.4% |
| rs4680 | *COMT* | 50%–75% reduction of the enzymatic activity [17] | Valine | Methionine | 33.8% |
| rs1800497 | *DRD2* | 12% less availability of striatal DRD2 [18] | A2 | A1 | 35.3% |

<sup>a</sup>MAFs (minor allele frequencies) were estimated in all of the study subjects (Suppl. Table S1).

Our case-control study included 361 schizophrenic patients and 282 healthy controls (Suppl. Table S1). None of the included SNPs showed an allelic or genotypic association with schizophrenia (Suppl. Table S2). However, we found that the individuals with the triple combination were significantly associated with schizophrenia (odds ratio [OR] 5.56, 95% confidence interval [CI] 1.25–24.69, p=0.011), whereas the double combination showed a tendency to be associated with schizophrenia (OR 2.54, 95%CI 1.00–6.46, p=0.062) (Tables 2, 3). Within the total of 16 individuals who carried the triple combination, 14 had been diagnosed as having schizophrenia.

**Table 2.** The double combination of dopamine-related functional SNPs

| Double combination | Schizophrenia patients |  | Healthy controls |  | Total |  |
| --- | --- | --- | --- | --- | --- | --- |
|  | n | Proportion | n | Proportion | n | Proportion |
| Candidate <sup>a</sup> | 19 | 5.4% | 6 | 2.2% | 25 | 4.0% |
| The rest | 336 | 94.6% | 270 | 97.8% | 606 | 96.0% |
| Total | 355 | 100.0% | 276 | 100.0% | 631 | 100.0% |

The association of the candidate combination was evaluated by Fisher's exact test.
<sup>a</sup>— The candidate double combination was T(+) of rs10770141 and Met(–) of rs4680, which showed a tendency to be related to schizophrenia (OR 2.54, 95%CI 1.00–6.46, $p=0.062$ ). The missing data: Six (1.7%) schizophrenia patients, six (2.1%) healthy controls; thus 12 (1.9%) for the total.

**Table 3.**
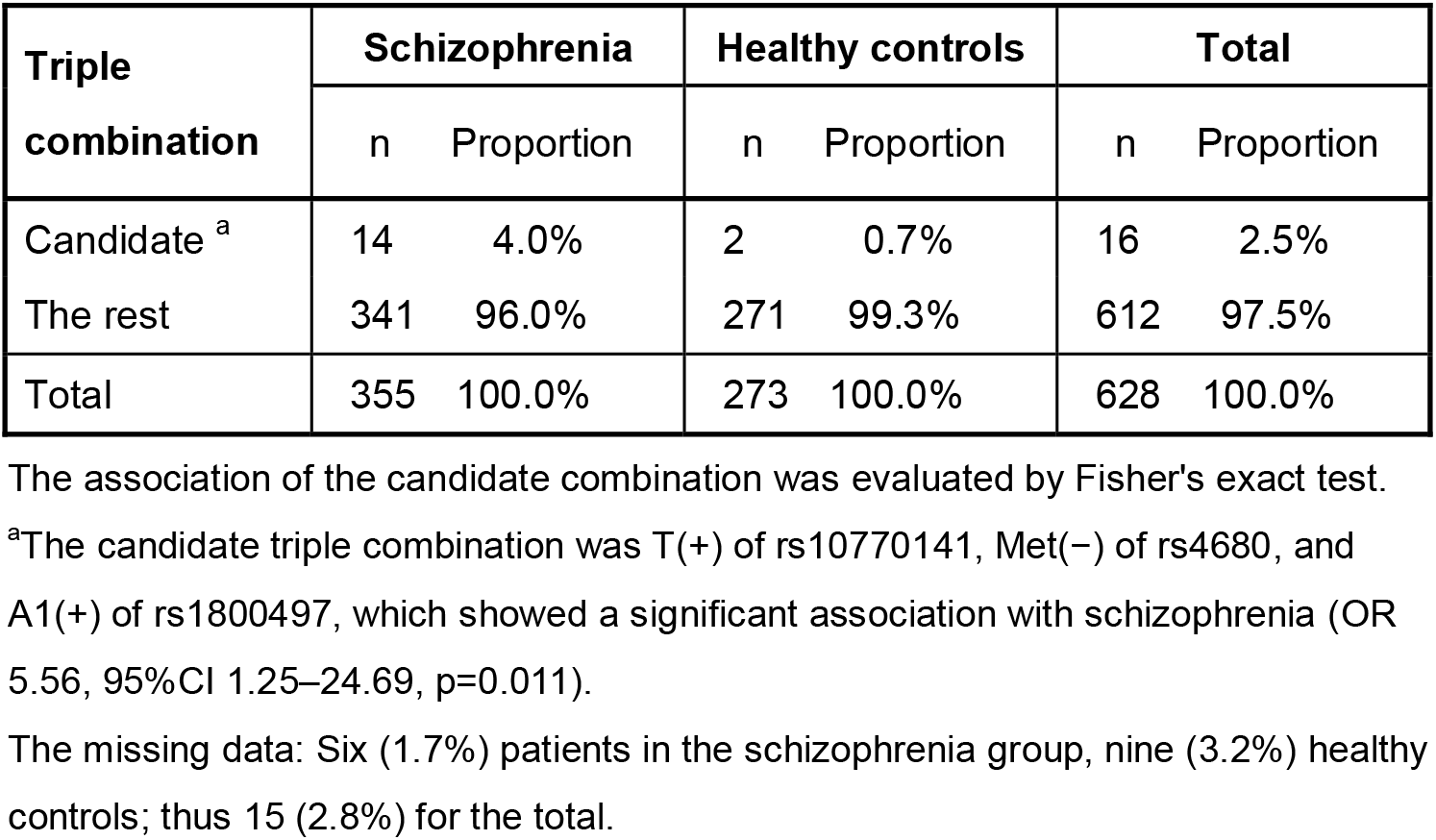
The triple combination of dopamine-related functional SNPs

| Triple combination | Schizophrenia |  | Healthy controls |  | Total |  |
| --- | --- | --- | --- | --- | --- | --- |
|  | n | Proportion | n | Proportion | n | Proportion |
| Candidate <sup>a</sup> | 14 | 4.0% | 2 | 0.7% | 16 | 2.5% |
| The rest | 341 | 96.0% | 271 | 99.3% | 612 | 97.5% |
| Total | 355 | 100.0% | 273 | 100.0% | 628 | 100.0% |

The association of the candidate combination was evaluated by Fisher's exact test.
<sup>a</sup>The candidate triple combination was T(+) of rs10770141, Met(–) of rs4680, and A1(+) of rs1800497, which showed a significant association with schizophrenia (OR 5.56, 95%CI 1.25–24.69, $p=0.011$ ).
The missing data: Six (1.7%) patients in the schizophrenia group, nine (3.2%) healthy controls; thus 15 (2.8%) for the total.

Although we observed that none of the single SNPs was associated with schizophrenia, a finding that is consistent with most of the literature, the significant association of the unique triple combination might indicate that individuals with schizophrenia have some inborn dopaminergic properties. Individuals with T(+) and Met(−) might be expected to show relatively higher dopamine synthesis and more rapid dopamine degradation, which also suggests that there are more dynamic alterations in dopamine concentrations at the synaptic cleft. In addition, carrying A1(+) might be associated with a lower presynaptic DRD2 density and consequently an enhanced dopamine release due to the attenuated negative feedback loops.^19^ Thus, these individuals might exhibit relatively more pulsatile dopamine signaling.

It is known that repeated exposure to dopamine agonists (such as psychostimulants^20^) and/or stressors^21^ causes sensitization. The term ‘sensitization’ means a progressive and long-lasting amplification of the behavioral and neurochemical response^22^ and has been argued to play a pathogenic role in causing not only psychosis but also schizophrenia.^23–25^ In fact, it is clinically suggested that frequent exposure to stress poses a risk for prodromal symptoms and schizophrenia.^26,27^ Individuals with schizophrenia were reported to release greater amounts of dopamine in response to stress.^6^ Individuals who carry the risk genetic combination might demonstrate similar dopaminergic pathogenesis, since relatively rapid pulsatile signaling could be caused repeatedly in response to daily stressors. This similarity in dopaminergic pathogenesis could contribute to sensitization and be a reason for the increased likelihood of schizophrenia onset. It may also be well consistent with the observations that both genetic and environment factors seem to present risks for schizophrenia.^28,29^

Our pathological hypothesis of schizophrenia demonstrated in this report accurately accounts for the recently proposed clinical picture of schizophrenia.^6,25^ Assuming that the sensitization to internal dopamine is somehow involved in the pathogenic mechanism of schizophrenia, the identification of the at-risk individuals at a preclinical stage might provide psychiatrists a pivotal chance to prevent the onset by managing the individual’s pulsatile dopamine activities. The risk genotypic combination discovered in the present study were detected in only 4.0% of the schizophrenic patients. However, the occurrence rate of the disease within the risk population was 87.5%, where the 14 out of 16 individuals were diagnosed to schizophrenia. Furthermore, although the polygenic risk score was reported to show no association with TRS^31^, the 9 out of these 14 schizophrenic patients (64.3%) were classified as TRS (Suppl. Table S3). We speculate that the inclusion rate could be improved by evaluating better combinations of other functional SNPs related to the dopamine pathway.

In addition, although the lowest MAF (5.7%, of rs10770141) was the apparently narrowing polymorphism in our study population, the Trans-Omics for Precision Medicine project recently reported to the U.S. National Center for Biotechnology Information that the MAF was estimated to be as large as 42.9% in the global population, suggesting its applicability.^30^ In the present study, we focused on the genetic variants related to the dopaminergic system, but our approach will be appropriate to assess other candidates related to different neurotransmissions including gamma-aminobutyric acid (GABA) and glutamatergic signaling.

There are some issues to be considered as study limitations. First, the mean age of our healthy control subjects was significantly lower than that of the patients. Although schizophrenia generally emerges during adolescence, our controls could still include potent false negatives. However, the impact may be limited because (1) the mean age of disease onset was significantly younger than the mean age of the controls, and (2) the lifetime prevalence of schizophrenia was estimated to be as small as 0.4%–1.0%. In addition, our sample size was large enough to allow us to evaluate our results, but further validations are clearly expected□—□ including those obtained for large and multi-ethnic populations.

In conclusion, we conducted a case-control study to investigate the association of functional SNPs related to dopamine transmission with schizophrenia. Our analyses revealed that the combination of the SNPs related to *TH, COMT* and *DRD2* genes was significantly associated with the onset of schizophrenia. We hypothesize that the present genetic combination characterizes dopamine transmission. The findings are important because of their potential to bring new insights to risk prediction and the development of new treatment methods for schizophrenia. However, future studies are necessary to test our findings.

All the authors have no conflicts of interests to disclose and approved this submission.

