## Supplementary material for "Genetic combination risk for schizophrenia": Methods

*Subjects*

This case-control study included the 361 patients who met the DSM IV-TR criteria for schizophrenia or schizoaffective disorder and 282 healthy controls. All of the participants were Japanese, and the patients were those who consulted Chiba University Hospital or affiliated hospitals in the period from May 2001 to October 2014. The healthy controls were recruited from the citizens and/or co-workers of the authors. The study was approved by the Ethics Committee of the Chiba University Graduate School of Medicine and was performed in accord with the Helsinki Declaration. The capability of patients to give informed consent was evaluated by the treating psychiatrist. Any patients who lacked adequate understanding of the study were excluded.

*Single nucleotide polymorphisms and genotyping*

We identified three SNPs to examine: rs10770141, rs4680 and rs1800497. Whole blood samples were obtained from all of the participants. The QIAamp DNA Blood Mini kit (250) (Qiagen, Valencia, CA) was used to isolate genomic DNA. Genotypes were determined by using TaqMan probe assays (Applied Biosystems, Foster City, CA) and the ABI PRISM 7300 Sequence Detection System (Applied Biosystems). In order to perform all of the procedures, we followed the protocols described in the manufacturer’s manuals. All polymerase chain reactions were conducted by one cycle at 95°C for 10 min, followed by 40 cycles at 92°C for 15 sec and 60°C for 60 sec.

*Quality assessments*

The quality of the study was evaluated by following the checklist of the Strengthening the Reporting of Genetic Association (STREGA) statement.^25^

*Statistical analysis*

Hardy-Weinberg equilibrium testing was used as a quality control for genotyping. Fisher's exact test, crude odds ratios (ORs), and 95% confidence intervals (CIs) were used to compare the risk genetic combinations model between groups. All statistical analyses were carried out using the Statistical Analysis System (SAS) software ver. 9.4 (SAS, Cary, NC). Statistical significance was set at p<0.05.
