## Supplemental Table 1 for "Genetic combination risk for schizophrenia"

**Suppl. Table S1.** The demographic data of the study subjects

|  | **Schizophrenia**  **n=361** | **Healthy controls**  **n=282** | | **p-value** |
| --- | --- | --- | --- | --- |
| Males / females | 178 / 183 | 134 / 143 | | 0.873^a^ |
| Age at evaluation, mean (SD), yrs | 50.5 15.4) | 38.8 (16.5) | | 0.000^b^ |
| Age at onset, mean (SD), yrs | 25.0 (8.3) |  |  |  |
| Age range, years | 16–92 | 14–85 | |  |

^a^Comparisons of two groups using chi-squared test.

^b^Comparisons of two groups using Student’s *t*-test.

The missing data: [sex] 5 (1.8%) for healthy control, then 4 (0.8%) for total; [age of evaluation] 4 (1.1%) for schizophrenia, 1 (0.4%) for healthy control, then 5 (0.8%) for total; [age of onset] 18 (5.0%) for schizophrenia.
