## Supplemental Table 2 for "Genetic combination risk for schizophrenia"

**Suppl. Table S2.** Genotypic distributions

| **SNP ID** | **Subjects** | **Genotype, n** | | | **Genotype, %** | | | **p-value ^a^** | **MAF, %** |
| --- | --- | --- | --- | --- | --- | --- | --- | --- | --- |
| ***TH gene*** | | | | | | | | | |
| rs10770141 | Schizophrenia (n=361)  Control (n=282) | T/T | T/C | C/C | T/T | T/C | C/C | 0.248 | 6.4 |
|  |  | 0 | 42 | 319 | 0 | 11.6 | 88.4 |  |  |
|  |  | 2 | 36 | 244 | 0.7 | 12.8 | 86.5 |  |  |
| ***COMT gene*** | | | | | | | | | |
| rs4680 | Schizophrenia (n=355)  Control (n=279) | Met/Met | Val/Met | Val/Val | Met/Met | Val/Met | Val/Val | 0.465 | 33.8 |
|  |  | 40 | 150 | 165 | 11.3 | 42.3 | 46.5 |  |  |
|  |  | 35 | 128 | 116 | 12.5 | 45.9 | 41.6 |  |  |
| ***D2R gene*** | | | | | | | | | |
| rs1800497 | Schizophrenia (n=361)  Control (n=278) | A1/A1 | A1/A2 | A2/A2 | A1/A1 | A1/A2 | A2/A2 | 0.370 | 35.3 |
|  |  | 52 | 159 | 150 | 14.4 | 44.0 | 41.6 |  |  |
|  |  | 40 | 108 | 130 | 14.4 | 38.8 | 46.8 |  |  |

^a^The p-values were obtained by chi-square tests. The missing data (*COMT* gene): six (1.7%) schizophrenia patients, three (1.1%) healthy controls; thus, 12 (1.9%) for the total. The missing data (*DRD2* gene): four (1.4%) healthy controls; thus four (0.6%) for the total. MAF, minor allele frequency; Met, methionine; SNP, single nucleotide polymorphism.
