## Supplemental Table 3 for "Genetic combination risk for schizophrenia"

**Suppl. Table S3.** The triple combination for comparing TRS

| **Triple**  **combination** | **TRS**  **patients** | **Non-TRS**  **patients** | **p-value** |
| --- | --- | --- | --- |
|  | n Proportion | n Proportion |  |
| Candidate ^a^ | 9 6.9%  121 93.1% | 5 2.2% | 0.0441 |
| The rest |  | 220 97.8% |  |
| Total | 130 100.0% | 225 100.0% |  |

The association of the candidate combination was evaluated by Fisher's exact test.

^a^ The candidate triple combination was T(+) of rs10770141, Met(−) of rs4680, and A1(+) of rs1800497.
